# Selection on heritable heterozygosity but no response to selection. Why?

**DOI:** 10.1101/026492

**Authors:** Tim Coulson, Sonya Clegg

## Abstract

The realisation that heterozygosity can be heritable has recently generated some elegant research. However, none of this work has discussed the fact that when heterozygote advantage occurs, heterozygosity can be heritable, yet allele frequencies remain at equilibrium and do not evolve with time. From a quantitative genetic perspective this means the character is heritable, is under selection, yet no response to selection is observed. We explain why this is the case, and discuss potential implications for the study of evolution in the wild.

## Main text

Heterozygosity-fitness correlations are often reported from natural populations (Hansson and Westerberg 2002; Chapman et al. 2009). Examples of heterozygosity averaged across multiple loci influencing life history traits are widespread (Hansson and Westerberg 2002), while sickle cell anaemia provides a text book example of heterozygote advantage at a single locus (Serjeant and Serjeant 1992). Heterozygosity can also be heritable, both at a single locus (Mitton et al. 1993; Nietlisbach and Hadfield 2015) and across multiple loci (Nietlisbach et al. 2015). Nietlisbach et al. (2015) point out that heritability of heterozygosity “is an essential requirement for an evolutionary response”. However, we would not always expect heterozygosity to evolve even when it is heritable and under selection. Here we explain why.

Heterozygote advantage maintains genetic variance: at a single locus, if the heterozygote is fitter than either homozygote, then genetic variation will be maintained, and allele frequencies will converge on equilibrium values (Crow et al. 1970). In the simplest case of two alleles and three genotypes *AA*, *AB* and *BB* we will denote the equilibrium allele frequencies of *A* as *p* and of *B* as *q* = 1 − *p*. When *p ≠ q* then heterozygosity at the locus will be heritable because the parent offspring covariance for heterozygosity will be greater zero (Mitton et al. 1993; Nietlisbach and Hadfield 2015). This means we have a case where heterozygosity is heritable, is under selection, but allele frequencies do not change and consequently no response to selection is observed. Why?

At a two allele locus the three genotypes *AA*, *AB* and *BB* occur at frequencies *p*^2^, 2*pq* and *q*^2^. They have relative fitnesses of 1 − *s*, 1 and 1 − *t*. The distribution of parental genotypes is consequently (1 − *s*)*p*^2^, 2*pq* and (1 − *t*)*q*^2^. However, because allele frequencies are at equilibrium, selection cannot alter them: *p* and *q* must take values such that the contrasting genotype frequencies before (*p*^2^, 2*pq* and *q*^2^) and after ((1 − *s*)*p*^2^, 2*pq* and (1 − *t*)*q*^2^) selection both have the same values of *p* and *q*. From a population genetics perspective this is well understood (Weir et al. 1990). Assume random mating. The genotype distribution among parents (((1 − *s*)*p*^2^, 2*pq* and (1 − *t*)*q*^2^2) must result in an offspring genotype distribution of *p*^2^, 2*pq* and *q*^2^ because genotype distributions cannot evolve indefinitely without shifting *p* and *q* – the equilibrium frequencies. So what does this mean from a quantitative genetic perspective, and in particular from the way that additive genetic variance is statistically estimated by comparing genotypes across relatives?

The narrow sense heritability is estimated as the phenotypic covariance between offspring and their parents. This, along with the heritability of a genotype or character, can be estimated using parent-offspring regression (Falconer 1960). Mitton et al. (1993) and Nietlisbach and Hadfield (2015) use population and quantitative genetic theory to explain the circumstances under which heterozygosity is heritable, and show how it can contribute to the additive genetic variance at a locus. This is elegant work. However, it is insufficient to explain the paradox we raise earlier: heterozygous advantage results in equilibrium allele frequencies even when heterozygosity is heritable. We show why by fitting parent-offspring regressions to genotype data at a locus when heterozygosity is heritable and subject to selection.

First, we denote the three genotypes -1, 0 and 1 rather than *AA*, *AB* and *BB*. This is because we need numbers in order to conduct a regression. We then regress these values in parents against those in offspring, weighting each point by the frequency of each genotype that each parental genotype will produce. For example, if we denote the genotype frequencies among selected parents as *f*_*AA*_, *f*_*AB*_ and *f*_*BB*_, then an *AA* parent is expected to produce *f*(*AA*)^2^ + 0.5*f*_*AA*_*f*_*AB*_ *AA* offspring. As is well understood, the regression line will have a slope of 0.5, which is interpreted as the heritability of the genotype (Figure 1A) (Weir et al. 1990). If we pool data across the two homozygotes, and now score homozygosity with a -1 and heterozygosity with a +1 we can conduct a similar analysis to estimate the heritability of heterozygosity (Figure 1B). As expected, when *p* ≠*q* then heterozygosity is heritable (Figure 1B), yet when *p* = *q* it is not (Figure 1C and 1D).

An important features of the regression lines in Figure 1 is that the intercept is not 0. In the example in Figure 1A, where *p* = 0.675 and *q* = 0.325 generated by setting *s* = 0.15 and *t* = 0.85 the regression line is *offspring genotype* = −0.175 + 0.5*parental genotype*, and in Figure 1B *offspring heterozygosity* = −0.1225 + 0.1225\**parent heterozygosity*.

The negative intercept arises because the key assumption of normality in parent-offspring regression is violated. The negative intercept is important here because it means that although offspring resemble their parents, selection cannot be realised because the inheritance process exactly counters the gains achieved by selection. Although we have only illustrated the negative intercept for a single locus with two alleles, a negative intercept will also be observed for multi-allelic loci and for multilocus genotypes at equilibrium. And this means that heterozygosity can not evolve with time when heterozygote advantage is operating and allele frequencies are at equilibrium, even though it will be heritable when *p ≠ q*.

A similar observation was observed by Coulson and Tuljapurkar (2008) when working with phenotypes in red deer – they noted that phenotypic inheritance exactly countered selection on birth weight because the intercept of the parent-offspring regression was less than zero. This was reflected by terms in the age-structured Price equation they used cancelling one another out. Coulson and Tuljapurkar (2008) did not propose a genetic mechanism for their observation; it is interesting to speculate that heterozygous advantage was the cause.

Not all loci are going to be at equilibrium, and when they are not, or when homozygote advantage is operating, then heterozygosity could evolve if it were subject to selection. However, it is unclear how frequently such circumstances will be encountered in the wild, because if selection is operating on a locus in the absence of heterozygote advantage we would expect allele frequencies to tend to fixation, although drift can obviously play a role in preventing this happening (Crow et al. 1970).

In many wild populations allele frequencies are not equal, heterozygosity is under selection, it is heritable and yet it is not evolving (Forcada and Hoffman 2014, for example). This is exactly what we would expect – heterozygosity may have a narrow sense heritability but in the broad sense it is not heritable. If heterozygosity contributes to a character’s phenotypic value it could also influence estimates of additive genetic variance and heritability of that character. Observations of heritable characters under selection that show no response to selection are widespread in natural systems (Merilä et al. 2001). Perhaps heritable heterozygosity provides an explanation in addition to those already proposed to explain this conundrum. Further theoretical and empirical work is required to see whether the non-additive component of the additive genetic variance is biasing estimates of heritability in the wild. In the meantime, it is probably worth while for quantitative geneticists to examine the intercept of parent-offspring regression as well as the slope.

## Acknowledgements

Thanks to the Centre for Advanced Study at the Norwegian Academy of Sciences who paid for the Heathrow-Oslo flight on which this manuscript was written.

## 1. Figure Legend

Figure 1. Parent offspring regressions on genotype (A) and (C) and heterozygosity (B) and (D) when *p* and *q* are not equal (A,B) and when they are (C,D). Regression lines are fitted through the data. Note that in (A) and (B) the parent-offspring regressions do not pass through (0,0).

## Footnotes

This manuscript was prepared with the AAS LATEX macros v5.2.

